# Repulsion of CA3 / dentate gyrus representations is driven by distinct internal beliefs in the face of ambiguous sensory input

**DOI:** 10.1101/2024.10.23.619862

**Authors:** Guo Wanjia, Subin Han, Brice A. Kuhl

## Abstract

Recent human neuroimaging studies of episodic memory have revealed a counterintuitive phenomenon in the hippocampus: when events are highly similar, corresponding hippocampal activity patterns are sometimes less correlated than activity patterns associated with unrelated events. This phenomenon—*repulsion—*is not accounted for by most theories of the hippocampus, and the conditions that trigger repulsion remain poorly understood. Here, we used a spatial route-learning task and high-resolution fMRI in humans to test whether hippocampal repulsion is fundamentally driven by internal beliefs about the environment. By precisely measuring participants’ internal beliefs and actively manipulating them, we show that repulsion selectively occurred in hippocampal subfields CA3 and dentate gyrus when visual input was ambiguous—or even *identical*—but internal beliefs were distinct. These findings firmly establish conditions that elicit repulsion and have broad relevance to theories of hippocampal function and to the fields of human episodic memory and rodent spatial navigation.

## INTRODUCTION

Central to the hippocampus’ role in episodic memory is its capacity to encode highly overlapping events while limiting potential interference^1–5^. Several recent human neuroimaging studies have identified a surprising way in which the hippocampus supports this goal: by inverting the representational structure of visual stimuli. That is, the hippocampus will—at least in some situations—form representations of overlapping events that are *less similar* (i.e., less correlated activity patterns) than representations of completely unrelated (non-overlapping) events^6–14^. This phenomenon has been termed *repulsion* because hippocampal representations of overlapping events appear to ‘move away’ from each other. While this phenomenon has now been observed several times, it is not entirely clear when or why repulsion occurs. Understanding the circumstances that elicit repulsion is of broad relevance to theories of memory and spatial navigation—both in humans and rodents—as leading theoretical accounts of the hippocampus largely fail to address or explain this surprising phenomenon^1,3,4,15^.

It is important to first note that repulsion is computationally distinct from traditional views of pattern separation^16^. The dominant view of pattern separation is that the hippocampus *orthogonalizes* input from sensory areas (e.g., input from visual cortex)^1,2,17^. A helpful way to conceptualize this is via an input-output function where, for every unit increase in the similarity of sensory input, the increase in hippocampal similarity (output) will be relatively smaller. With perfect orthogonalization, increases in input similarity would not increase similarity in the hippocampus at all. In other words, the ceiling for orthogonalization is a flat input-output function. In contrast, repulsion occurs if an increase in input similarity leads to a *decrease* in hippocampal similarity—i.e., a portion of the input-output function that is *negatively sloped*.

To understand why hippocampal repulsion occurs, it is essential to understand when it occurs. While event similarity is, by definition, a necessary ingredient for repulsion, similarity does not always—or automatically—induce hippocampal repulsion. Indeed, there are many examples where hippocampal pattern similarity is relatively greater when events have overlapping elements^18–21^. Based on current evidence, one factor that seems to be important for inducing repulsion is the degree of experience with overlapping events. Namely, repulsion may only emerge with extensive training^6,16^. However, experience, per se, is not a satisfying explanation—rather, experience is presumably correlated with some change in behavior or memory that explains why hippocampal representations ultimately exhibit repulsion^9^.

Perhaps the most intuitive explanation is that repulsion only emerges once participants learn to visually attend to subtle differences between otherwise similar stimuli. However, a visual attention account predicts that effects should first emerge in visual cortical areas and only then be passed on to the hippocampus. In contrast, repulsion effects in the hippocampus have been shown to occur without any evidence of precipitating effects in visual cortex^6,7,9^. That said, a modified version of this account could be that the hippocampus amplifies or exaggerates subtle differences in visual cortical areas.

An alternative account is that repulsion has less to do with differences in visual attention and more to do with differences in internal beliefs. From this perspective, the hippocampus is not inheriting or amplifying differences from visual cortex but is, instead, generating these differences internally. This account is motivated by recent theory^22^ and evidence from studies of rodents^23,24^ which argue that changes in hippocampal activity patterns (place cell remapping) are better explained by shifts in internal (or latent) representations than by observable features of the environment. That said, it is not clear whether theories of place cell remapping in rodents apply to the phenomenon of hippocampal repulsion in human memory. Indeed, the phenomenon of repulsion has not been reported in rodent place cells (however, it has been anticipated in computational models^25^).

Here, we sought to test whether hippocampal repulsion specifically occurs when internal beliefs are distinct, but visual stimuli are ambiguous. To this end, we used high-resolution fMRI and a carefully-designed spatial route-learning paradigm in which human participants learned pairs of overlapping routes. Inspired by classic rodent T-maze designs^26,27^, the routes were constructed such that they initially overlapped but eventually diverged. More specifically, during the initial route segment, the overlapping route images were visually identical (‘same segment’); the routes then became very slightly different (‘similar segment’) before ultimately diverging (‘different segment’) and terminating at unique destinations. This allowed us to assess the similarity of fMRI activity patterns—in the hippocampus and visual cortical areas—as a function of route segment. Additionally, we took two approaches to linking hippocampal activity patterns to internal beliefs. First, we obtained participant-specific measures of the timepoint within each route when participants were able to confidently predict a route’s destination, allowing us to test whether hippocampal repulsion was temporally-coupled with these ‘moments of insight.’ Second, we directly manipulated beliefs by using probabilistic cues that indicated the likely route destination. This afforded a causal test of whether distinct beliefs, in the face of ambiguous input, drive hippocampal repulsion effects.

To preview, we show that hippocampal subfields CA3 and dentate gyrus (CA3/DG) exhibited repulsion effects when visual input was extremely similar—or even *identical*—but internal beliefs were distinct. These findings not only provide insight into when and why repulsion occurs, but they establish the relevance of this poorly-understood representational phenomenon to memory interference and spatial navigation.

## RESULTS

Each participant repeatedly viewed a slideshow of images depicting four routes (2 overlapping route pairs) from the University of Oregon campus. Within each overlapping pair, the two routes started on identical paths with identical images (‘same segment’, 6s, 25 pictures), followed by identical paths with highly similar images (‘similar segment’, 12s, 50 pictures), then different paths with distinct images (‘different segment’, 6s, 25 pictures) (**Fig 1A**). Each route terminated at a unique landmark, or ‘destination’, which was identified by a text label (e.g., ‘pole’) that appeared on screen for 2s. Participants gained initial familiarity with the four routes before fMRI scanning and then repeatedly viewed each route again during scanning. Critically, during fMRI scanning—and only during fMRI scanning—each trial was preceded by a probabilistic cue (75% valid) indicating the likely destination. When invalid, the cue always indicated the destination of the overlapping route.

**Fig 1.**
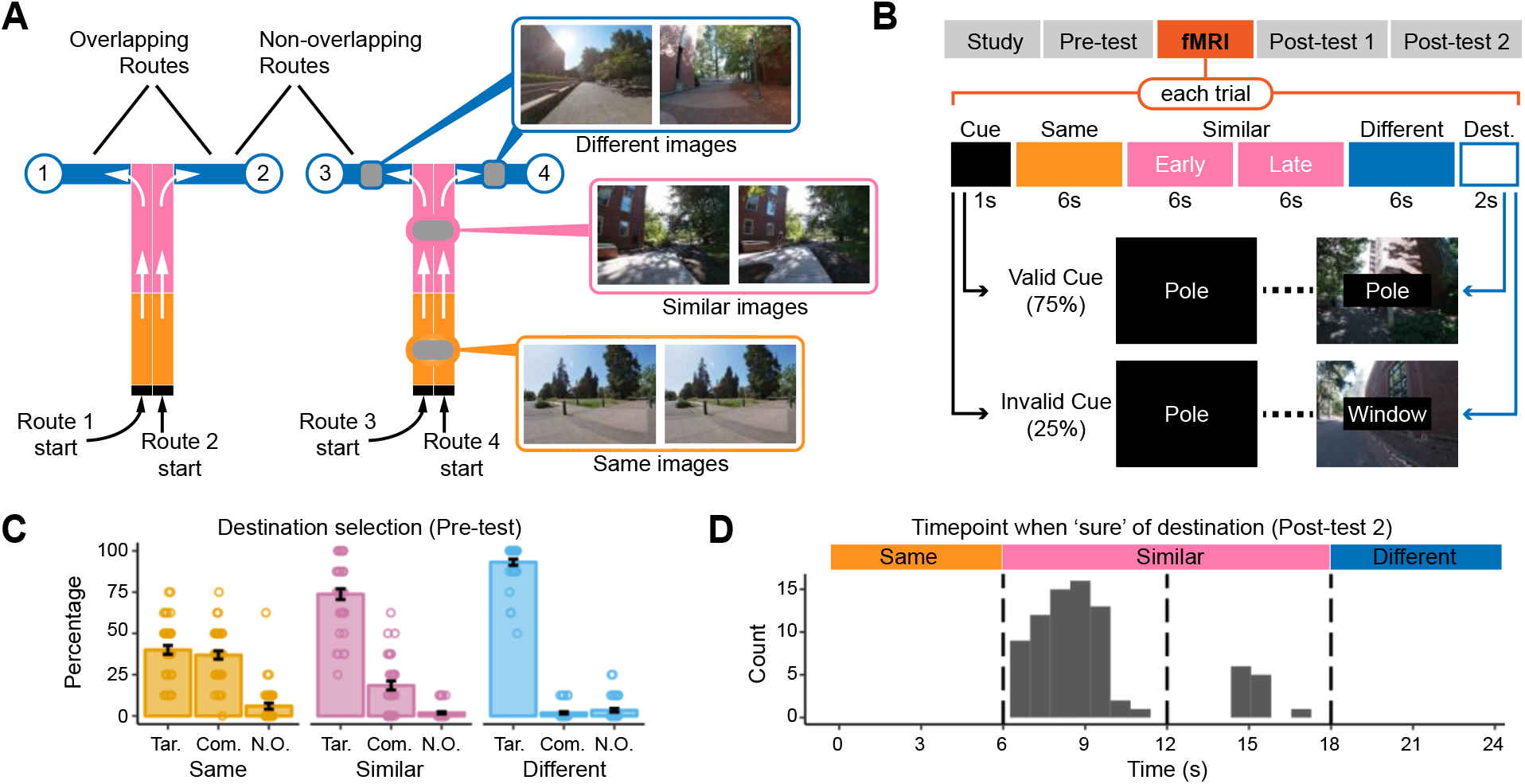
Experimental paradigm and behavioral results. (A) Schematic illustration of overlapping and non-overlapping routes. Each participant studied two pairs of overlapping routes. Overlapping routes (e.g. routes 1 and 2) initially followed identical paths with identical images (‘same segment’, orange) before continuing on identical paths but with subtly different images (‘similar segment’, pink) and then diverging on different paths with different images (‘different segment’, blue) to terminate at unique destinations (circles). For non-overlapping routes (e.g., routes 1 and 3), paths and images were always distinct. **(B)** Overview of experimental phases and fMRI trials. During the fMRI phase, each trial was preceded by a cue (1s) indicating the likely destination (75% valid; 25% invalid). Invalid cues always referred to the overlapping route’s destination. The cue was followed by a stream of 100 images displayed across 24s (same segment = 6s; similar segment = 12s; different segment = 6s). The destination was then displayed for 2s. **(C)** During the Pre-test, routes paused during each segment and participants were instructed to choose the correct destination. Within the same segment, participants were equally likely to choose the target destination (Tar.) or competitor destination (Com.) (p = 0.447) and chose both more often than a non-overlapping route’s destination (N.O.) (p’s < 0.001). Within the similar segment, the target was selected more often than competitor or non-overlapping destinations (p’s < 0.001), and the competitor more often than a non-overlapping destination (p < 0.001). Within the different segment, the target was selected more often than the competitor destination (p < .001), but competitor and non-overlapping destinations were equally likely to be selected (p = .201). **(D)** In Post-test 2, participants viewed routes and were instructed to press a button when they were ‘90% sure’ of the destination. The histogram shows the distribution of responses across time. Note: error bars represent S.E.M.

### Behavioral Results

Before, during and after scanning, participants were tested, in several different ways, on their ability to predict where each route’s destination. Prior to fMRI scanning, participants completed two rounds of a Pre-Test in which the route slideshow paused and participants were asked to select the route destination from the set of all 4 possible destinations. After making a selection, participants indicated whether their confidence was high or low. Pauses occurred equally often during the same, similar, and different segments, allowing for consideration of performance at each segment. Throughout the manuscript, we refer to the correct destination as the ‘target,’ the destination corresponding to the overlapping route as the ‘competitor,’ and the two destinations from non-overlapping routes as the ‘non-overlapping’ destinations. The key measure of interference resolution was the ability to select the target destination over the competitor destination. Note: the probability of selecting the target by chance was 25%, the probability of selecting the competitor was 25%, and the probability of selecting a non-overlapping destination was 50%.

As expected, performance in the Pre-Test improved as a function of route segment (see **Supplementary Table 1** and **Fig. 1C**). During the same segment (where the images were identical for the overlapping routes), participants were more likely to select the target or competitor destination than a non-overlapping destination (target vs. non-overlapping: t_39_ = 8.62, p < 0.001; competitor vs. non-overlapping: t_39_ = 8.85, p < 0.001), but, as expected, did not differ in the likelihood of selecting the target vs. competitor (t_39_ = 0.77, p = 0.447). During the similar segment, participants were more likely to select the target than either the competitor (t_39_ = 9.93, p < 0.001) or a non-overlapping destination (t_39_ = 20.60, p < 0.001). However, participants were still more likely to select the competitor than a non-overlapping destination (t_39_ = 5.90, p < 0.001). Finally, during the different segment, participants were again more likely to select the target than the competitor (t_39_ = 39.83, p < 0.001) or a non-overlapping destination (t_39_ = 32.56, p < 0.001), but, importantly, they were no more likely to select the competitor than a non-overlapping destination (t_39_ = 1.30, p = 0.201). Thus, competition between the overlapping routes was highest during the same segment (when overlapping routes had identical images), was lower but still present during the similar segment, and was fully resolved during the different segment. The percentage of trials on which the target was selected with high-confidence also robustly increased from the same segment to the similar segment (t_39_ = 12.76, p < 0.001) and from the similar segment to the different segment (t_39_ = 9.41, p < 0.001) (**Supplementary Table 1**).

During fMRI scanning, routes occasionally paused (25% of trials) and participants were instructed to select the correct destination. However, these trials were only included for the sake of promoting participant engagement and vigilance; performance on these trials is not easily interpreted given that they were always preceded by a valid cue (see Methods).

After fMRI scanning, participants completed two Post-tests. The first Post-test was similar to the Pre-Test except that the routes only paused during the similar segment and only the target and competitor destinations were shown. Each route was tested repeatedly, with pauses occurring at picture indices 30, 45, 60, and 75. A one-way repeated measures ANOVA revealed a significant main effect of picture index on the probability of selecting the target (F_3, 117_ = 88.98, p < 0.001) and on the percentage of trials for which the target was selected with high confidence (F_3, 117_ = 164.07, p < 0.001; see **Supplementary Table 2**). Splitting the similar segment into an early-similar segment (picture index 30 and 45) and late-similar segment (picture index 60 and 75) revealed a significant increase, from early-similar to late-similar, in target selection (t_39_ = 8.5217, p < 0.001) and in target selection with high confidence (t_39_ = 12.25, p < 0.001).

In the second Post-test, participants viewed the routes again and were instructed to press a button as soon as they were at least ‘90% sure’ of the destination. Upon making a button press, the route paused and participants were then prompted to select the destination from the full set of 4 destinations. Participants selected the target on 94.84 (mean) ± 7.99% (standard deviation) of the trials. For trials on which the target was selected, the mean response time was 9.36 ± 1.96s (relative to the start of the route), which corresponded to the first half of the similar segment (early-similar segment). While there was variability across participants (min: 6.99s; max: 12.77s; **Fig. 1D**), all participants had a mean response time that fell somewhere within the similar segment (which spanned 6.0 to 18.0s).

### fMRI Pattern Similarity as a Function of Route Segment

For each participant, trial, and region of interest (ROI), we extracted each voxel’s activation at each 1s timepoint, excluding the cue and destination (thus, 24 timepoints were included in total). Following prior studies^6,7,9^, we focused analyses on the hippocampus, parahippocampal place area (PPA) and early visual cortex (EVC). We specifically considered two sub-regions of the hippocampus: CA1 and CA3/dentate gyrus (CA3/DG) (**Fig 2A**, see Methods). We predicted repulsion effects would be selective to CA3/DG, based on findings from related studies^9,11,19,23^.

**Fig 2.**
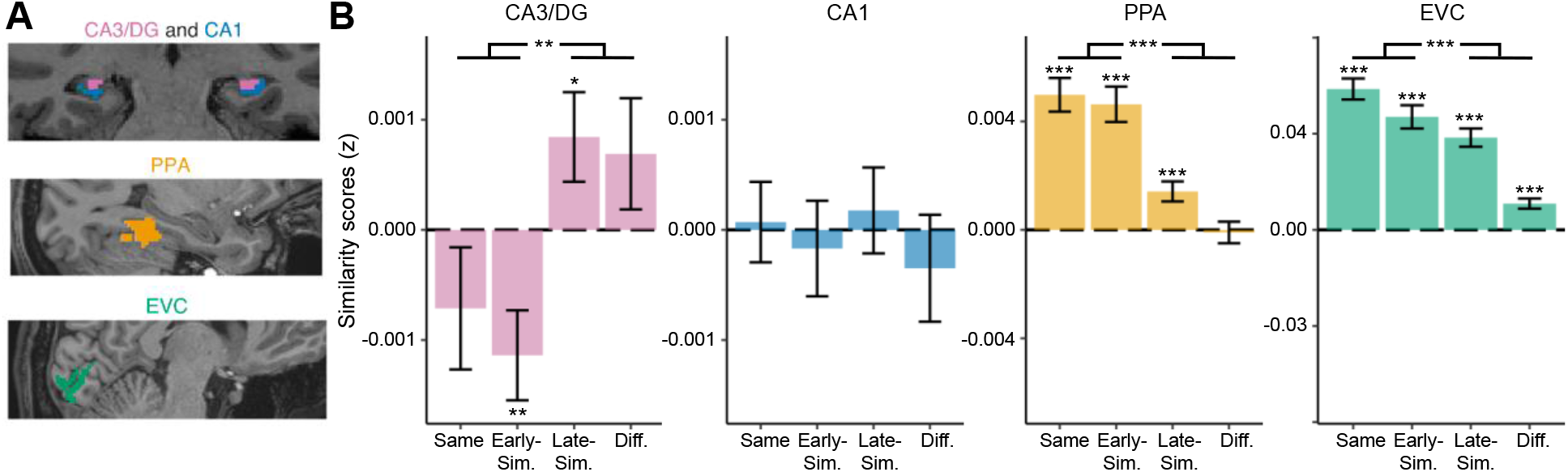
fMRI similarity scores as a function of route segment. (A) Regions of interest (ROIs) included hippocampal subfields CA3 and dentate gyrus (CA3/DG, pink), hippocampal subfield CA1 (blue), the parahippocampal place area (PPA, yellow), and early visual cortex (EVC, green). **(B)** Similarity scores were defined as the fMRI pattern similarity between overlapping routes relative to pattern similarity between non-overlapping routes. Positive similarity scores indicate that overlapping routes had more similar activity patterns than non-overlapping routes. Negative similarity scores indicate repulsion: that overlapping routes had less similar activity patterns than non-overlapping routes. Within CA3/DG, similarity scores significantly increased from the first half of the route to the second half (p = .001), with evidence of repulsion selective to the first half of the trial. In contrast, similarity scores significantly decreased from the first to second half of the trial in PPA and EVC (p’s < .001), but with no evidence of repulsion. Notes: error bars represent S.E.M.; * p < .05; ** p < .01; *** p < .001.

Pattern similarity values were computed, on a timepoint-by-timepoint basis, by correlating activity patterns from different routes across different scan runs. Correlations between non-overlapping routes were subtracted from correlations between overlapping routes, yielding a *similarity score*. Positive similarity scores indicate that overlapping routes were more similar than non-overlapping routes. Negative similarity scores indicate that overlapping routes were less similar than non-overlapping routes (evidence for repulsion). It is important to emphasize that overlapping and non-overlapping routes do not refer to different *routes*^28^, but to different *comparisons* between routes. For example, whereas routes 1 and 2 would be considered overlapping routes, routes 2 and 3 would be non-overlapping routes (**Fig. 1A**). For initial analyses, we only included trials that were preceded by valid destination cues (invalid cues are considered later) and we divided each route into four evenly-spaced segments (6s each): same, early-similar, late-similar, different. While overlapping routes were not discriminable based on visual information during the same segment, the valid destination cues allowed participants to ‘believe’ that the routes were distinct even during the same segment.

A 2-way ANOVA with factors of segment and ROI (CA3/DG, CA1, PPA, EVC) revealed significant main effects of segment (F_3,585_ = 33.74, p < 0.001) and ROI (F_3,585_ = 375.76, p < 0.001), as well as a significant interaction (F_9,585_ = 26.10, p < 0.001). The interaction indicated that changes in similarity scores across segments differed across ROIs (**Fig. 2B**). Considering each ROI separately, significant main effects of segment were observed in CA3/DG (F_3,177_ = 5.15, p = 0.002), PPA (F_3,117_ = 28.71, p < 0.001), and EVC (F_3,117_ = 40.91, p < 0.001), but not in CA1 (F_3,177_ = 0.36, p = 0.783). Whereas similarity scores in PPA and EVC robustly decreased across segments (paired samples t-test of first half vs. second half of the trial; PPA: t_39_ = 7.21, p < 0.001; EVC: t_39_ = 8.64, p < 0.001), similarity scores in CA3/DG significantly *increased* across time bins (first half vs. second half: t_39_ = -3.44, p = 0.001). Thus, CA3/DG representations of overlapping routes were most distinct when the images were most similar. In fact, similarity scores in CA3/DG were significantly below 0 during the first half of the trial (same + early similar segment, one sample t-test: t_39_ = -2.30, p = 0.027). In other words, CA3/DG exhibited a repulsion effect (lower pattern similarity for overlapping routes than non-overlapping routes) that specifically occurred when the overlapping routes were identical or highly similar.

It is also notable that, within the similar segment alone, there were abrupt changes from the early-similar to late-similar segments in CA3/DG, PPA and EVC. Again, however, these changes were in opposite directions. Whereas PPA/EVC similarity scores sharply decreased from the early-similar to late-similar segments (paired samples t-tests; PPA: t_39_ = 4.77, p < 0.001; EVC: t_39_ = 2.51, p = 0.016), CA3/DG scores sharply *increased* from the early-similar to late-similar segments (t_39_ = -3.81, p < 0.001). In fact, CA3/DG scores were significantly below 0 in the early-similar segment (t_39_ =-2.79, p = 0.008) and significantly above 0 during the late-similar segment (t_39_ = 2.08, p = 0.045). The fact that similarity scores markedly changed within the similar segment is consistent with the behavioral data from Post-test 2, which indicated that the ability to discriminate between the target and competitor destination emerged, for each participant, during the similar segment (**Fig. 1D**).

### Relationship Between Repulsion and ‘Moments of Insight’

The findings described above support our prediction of a repulsion effect within CA3/DG. We next sought to establish a direct link between CA3/DG repulsion and internal beliefs about a route’s destination. We predicted that repulsion would be temporally-coupled to the specific moment *within a trial* when participants were able to confidently disambiguate the overlapping routes. To test this, we used each participant’s responses from Post-test 2 to identify the exact timepoint when participants were highly confident (‘90% sure’) of each route’s destination. We refer to this critical timepoint as the ‘Moment of Insight’ (MoI). Importantly, Post-test 2 did not include probabilistic destination cues; thus, the data provide a measure of when participants were able to use subtle visual cues *within the route images* to determine the route destination. As in the preceding section, here we focus only on fMRI data from valid trials.

To compute the MoI, for each participant we pooled the Post-test 2 responses within each route pair. Pooling responses within a pair was necessary because the fMRI similarity scores were computed at the level of route pairs, not individual routes. The MoI for each pair was defined as the range (minimum to maximum) of the pooled responses. The mean duration of the MoI (i.e., the difference between the minimum and maximum values) was 1.68s, indicating that the MoI was a precise (narrow) temporal window relative to the full 24s trial (**Fig. 3A**). Timepoints prior to the MoI were defined as pre-MoI and timepoints after the MoI as post-MoI (**Fig 3A**). The pre-MOI segment had an average length of 8.70s and the post-MoI segment had an average length of 13.63s.

**Fig 3.**
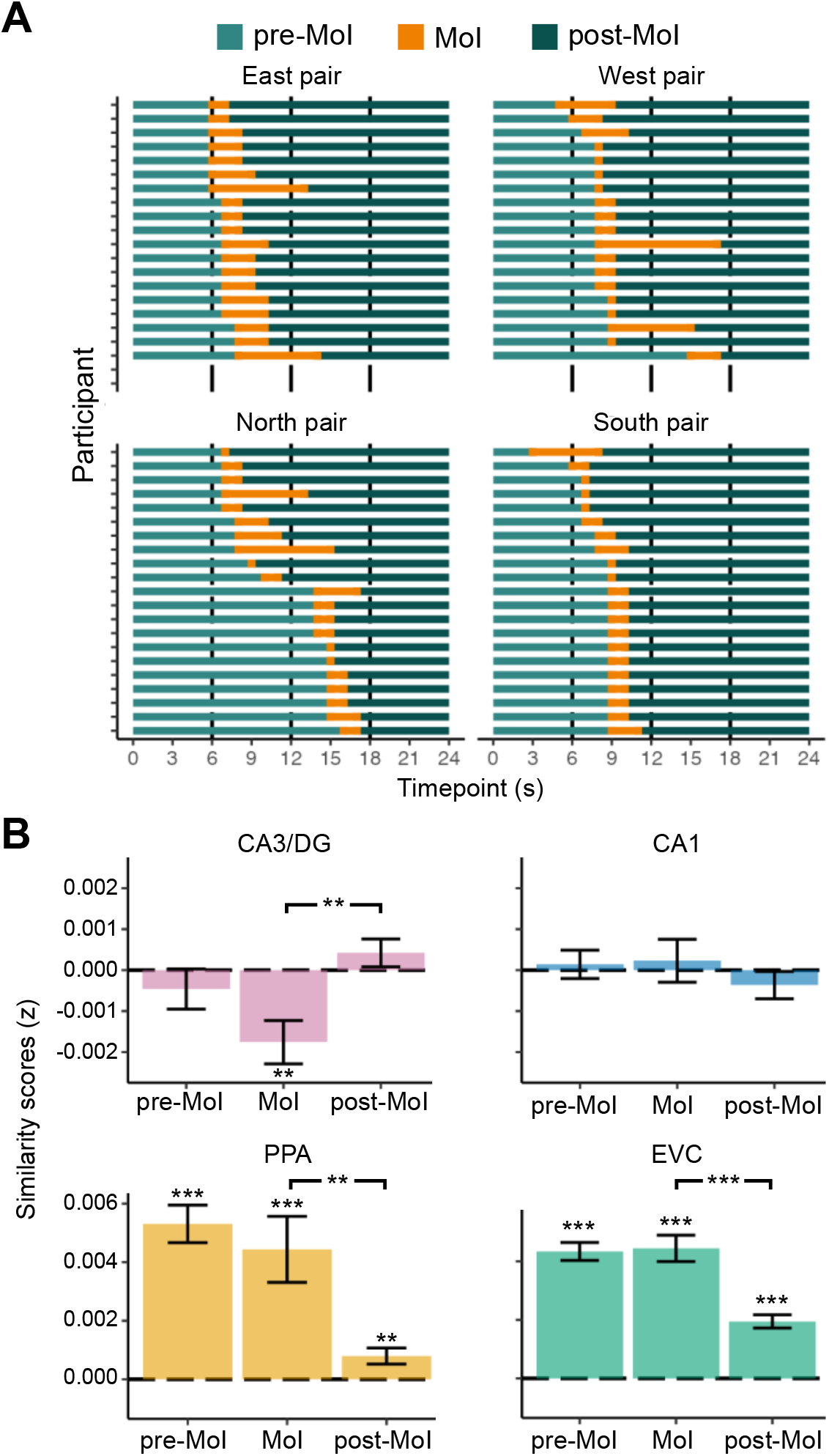
CA3/DG repulsion is temporally coupled to Moments of Insight. (A) Distribution of Moments of Insight (MoI) across route pairs and participants. Each participant studied two pairs of overlapping routes (east and west or south and north). For each participant and each pair of overlapping routes, the MoI was defined based on participants’ responses during Post-test 2 (responses indicated the moment at which participants were ‘90% sure’ of the destination). Within each of the four plots, each row corresponds to an individual participant. For visualization purposes, the rows (participants) are vertically rank-ordered within each plot by the timing of the MOI. The width of the MoI (orange bars) represents the range of responses (minimum to maximim). **(B)** fMRI similarity scores at participant-specific MoI, as well as pre-MoI and post-MoI segments. For CA3/DG, a significant repulsion effect (similarity scores < 0) was observed at the MoI (p = 0.002). Similarity scores in CA3/DG significantly increased from the MoI to post-MoI segment (p = 0.002). In PPA and EVC, similarity scores showed an opposite pattern: significant decreases from the MoI to post-MoI segments (PPA: p = 0.001; EVC: t_39_ = 7.64, p < 0.001). Notes: error bars represent S.E.M.; ** p < .01; *** p < .001.

CA3/DG similarity scores were robustly below 0 (repulsion effect) at the MoI (one-sample t-test; t_39_ = -3.32, p = 0.002; **Fig. 3B**), but did not differ from 0 pre-MOI (t_39_ = -0.95, p = 0.350) or post-MOI (t_39_ = 1.24, p = 0.222). Considering all three segments (pre-MOI, MOI, post-MOI), there was a significant main effect of segment in CA3/DG (one-way ANOVA: F_2,78_ = 5.74, p = 0.005). This was primarily driven by a significant increase in similarity scores from the MoI to post-MoI segment (paired samples t-test: t_39_ = 3.28, p = 0.002). There was no significant difference between pre-MoI and MoI (t_39_ = -1.75, p = 0.088). For CA1, EVC, and PPA, there was no evidence of repulsion effects at the MoI (all similarity scores numerically positive). The main effect of segment was significant for EVC (F_2,78_ = 30.79, p < 0.001) and PPA (F_2,78_ = 11.34, p < 0.001), but not CA1 (F_2,78_ = 0.69, p = 0.506). However, in sharp contrast to CA3/DG, the main effects for EVC and PPA were driven by significant *decreases* in similarity scores from the MoI to post-MoI segments (t_39_’s > 3.44, p’s < 0.001; **Fig. 3B**). Thus, after participants gained insight into where the routes were headed, representations of overlapping routes became much less similar in EVC and PPA, but much more similar in CA3/DG.

Finally, to further confirm the temporal selectivity of the CA3/DG repulsion effect to the MoI, we performed a permutation test. For this analysis, the similarity scores were computed for each participant, route pair, and timepoint within a route, as described above. These 24 timepoints were then shuffled (separately for each route pair and participant) before the MoI analysis proceeded. This process was repeated 10,000 times to obtain a distribution of permuted group-level mean similarity score values at the MoI. The actual (observed) mean similarity score at the MoI was lower than any of the 10,000 permuted means (i.e., p < .0001), strongly demonstrating that the repulsion effect at the MoI was stronger than what would be observed by randomly selecting timepoints within the routes. Taken together, these analyses demonstrate a repulsion effect that was highly selective to CA3/DG and clearly time-locked to the moment when participants were able to internally disambiguate the overlapping routes.

### Relationship Between Destination Cues and Repulsion

Because the moments of insight (MoI) were defined based on trials without any destination cues (Post-test 2), they were intended to capture internal beliefs *not explained* by the destination cues. In a final set of analyses, we considered a complementary question: whether the destination cues, on their own, were sufficient to drive CA3/DG repulsion effects. Given our overarching hypothesis that CA3/DG repulsion is driven by internal beliefs, we predicted that similarity scores in CA3/DG would be influenced by destination cues, even when fully controlling for sensory information.

In all of the fMRI analyses presented thus far, pattern similarity between overlapping routes was computed using pairs of trials that were preceded by valid cues (‘valid-valid’ similarity scores). Here, we contrast these ‘valid-valid’ similarity scores with ‘valid-invalid’ similarity scores (**Fig. 4A**). For the valid-invalid similarity scores, we again computed pattern similarity of overlapping routes relative to non-overlapping routes, but each pair of overlapping routes consisted of a validly-cued route and an invalidly-cued route. Thus, a trial where cue = X and destination = X (valid trial) would be correlated with a trial where cue = X and destination = Y (invalid trial). In terms of the actual route images that participants saw, there was absolutely no difference between a pair of valid-valid routes and a pair of valid-invalid routes. In both cases, the pair of routes had same, similar and different segments, ending at distinct destinations. The only difference was that for the valid-valid routes, participants (correctly) *expected* the routes to end at distinct destinations (X vs. Y), whereas for the valid-invalid routes, participants (incorrectly) expected the routes to end at the same destination (X vs. X). We predicted that any effects of destination cues (valid-valid vs. valid-invalid) on CA3/DG representations would be strongest during the early part of the trial—when visual information was most ambiguous.

**Figure 4.**
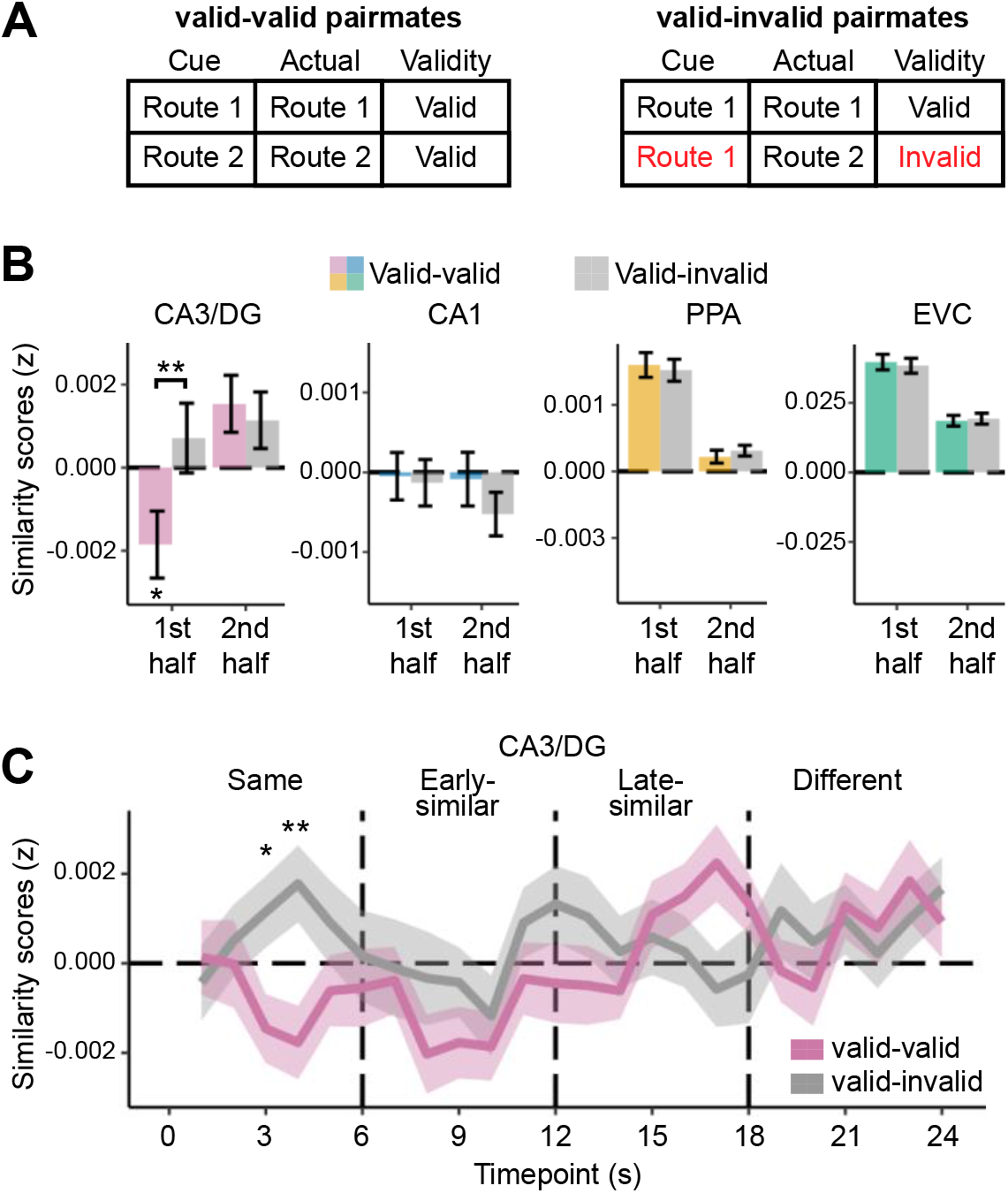
Influence of destination cues on similarity scores. (A) Schematic illustration of how valid-valid pairmates (left) and valid-invalid pairmates (right) were defined. In both cases, fMRI pattern similarity was computed between overlapping routes (e.g., route 1 and route 2). In the valid-valid comparison, the cues accurately indicated that the routes were headed to different destinations. However, in the valid-invalid comparison, the cues were the same (e.g., ‘route 1’) but the destinations were different. **(B)** fMRI similarity scores as a function of cues (valid-valid vs. valid-invalid) and route half (1^st^ half of the route = picture index 1-50 vs. 2^nd^ half of the route = picture index 51-100). Only CA3/DG showed a significant interaction between cues and route half (p = 0.038). For CA3/DG, within the first half of the route, valid-valid similarity scores were significantly below 0 (repulsion effect; p = 0.027) and significantly lower than valid-invalid similarity scores (p = 0.009). **(C)** Timepoint-by-timepoint similarity scores in CA3/DG, separated by valid-valid and valid-invalid comparisons. Within the same segment (when route images were identical), valid-valid similarity scores were significantly lower than valid-invalid similarity scores at timepoints 3 and 4. Notes: error bars represent S.E.M.; * p < .05; ** p < .01.

As a first step, we compared the first half of each route (same + early similar segments) vs. the second half (late-similar + different segments). For each ROI, we ran two-way ANOVAs with factors of half (1^st^, 2^nd^) and cue (valid-valid, valid-invalid). Within CA3/DG, there was a significant main effect of half (F_1,117_ = 7.31, p = 0.008), reflecting an overall increase in similarity scores from the first to second half, but no main effect of cue (F_1,117_ = 2.35, p = 0.128). However, there was a significant interaction between half and cue (F_1,117_ = 4.39, p = 0.038; **Fig. 4B**), driven by valid-valid similarity scores that were significantly lower than valid-invalid similarity scores in the first half of the trial (paired t-test: t_39_ = - 2.73, p = 0.009), but not in the second half of the trial (t_39_ = 0.45, p = 0.655). In EVC and PPA (but not CA1), there were significant main effects of half (PPA: F_1,117_ = 131.85, p < 0.001, EVC: F_1,117_ = 213.13, p < 0.001, CA1: F_1,117_ = 0.53, p = 0.469), reflecting an overall decrease in similarity scores, but there were no main effects of cue or interactions between cue and half (F_1,117_’s < 0.78, p’s > 0.380; **Fig. 4B**). When considering the first half of the trial alone, there was no difference between valid-valid and valid-invalid similarity scores for CA1, PPA, or EVC (t_39_’s < 2.01, p’s > 0.05). Thus, destination cues selectively influenced the relative similarity of competing route representations within CA3/DG, particularly when visual information was most ambiguous.

To fully isolate the influence of cues (and corresponding internal beliefs), we conducted an additional, timepoint-by-timepoint analysis that specifically focused on the 6 timepoints during the same segment—when overlapping route images were identical. Any effect of cues during the same segment can *only* be explained by the cues themselves, since the route images were identical across overlapping routes during this segment. Again, we compared valid-valid to valid-invalid trials. To account for multiple comparisons across the 6 timepoints, we used a Bonferroni-corrected threshold of p < .0083 (uncorrected) to establish significance (i.e., p = .05/6). Strikingly, valid-valid similarity scores were significantly lower than valid-invalid similarity scores at timepoint 4 (t_39_ = -3.32, p = 0.002; **Fig. 4C**), with a similar effect at timepoint 3, though not significant at the Bonferroni corrected threshold (t_39_ = -2.69, p = 0.010). Qualitatively, the valid-valid and valid-invalid comparisons were mirror-images of each other, with valid-valid similarity scores below 0 and valid-invalid similarity scores above 0. Thus, when viewing identical route images, repulsion occurred— but only occurred—when internal beliefs were distinct.

## DISCUSSION

Here, we show that representations of overlapping spatial routes in hippocampal subfields CA3 and dentate gyrus (CA3/DG) exhibit a dramatic repulsion effect that specifically occurs when visual input to the hippocampus is highly similar—or even *identical*—but internal beliefs are distinct. By using a human fMRI paradigm that was inspired by rodent studies^26,27^, and with predictions derived from properties of place cell remapping^22,23^, our approach and findings uniquely bridge the fields of human episodic memory and rodent spatial navigation. More generally, our findings are broadly relevant to theories of hippocampal function, which largely do not account for or anticipate the phenomenon of repulsion^1–4^.

While the phenomenon of repulsion in the human hippocampus has now been reported several times^6–14^, the current study provides a unique and theory-driven test of when and why repulsion occurs. Our prediction that repulsion would be linked to internal beliefs was inspired by recent theoretical and empirical arguments that place cell remapping in rodents is better explained by changes in internal beliefs than by external features of the environment^22,23^. That said, the phenomenon of repulsion has not been reported in rodent place cells. In humans, there is evidence that internal beliefs about ambiguous stimuli are related to hippocampal activity patterns^29,30^, but not specifically to the phenomenon of repulsion. There is also evidence that repulsion in the human hippocampus is correlated with behavioral expressions of memory interference^7,9,16^, but this evidence is potentially compatible with an internal belief account or an account based on selective attention.

Our first approach to linking hippocampal repulsion to internal beliefs involved using participant-specific subjective reports (collected after fMRI scanning) to identify the precise timepoint, within each route, when participants were ‘sure’ of the route’s destination (*moment of insight*; MoI). Because the MoI were based on participants’ beliefs in the absence of probabilistic cues, they reflected a participant’s ability to discern the route destination from subtle visual features of the routes. Notably, the MoI always occurred at some point during the similar segment—that is, while the route images were still extremely similar (**Fig 1**). Despite this high visual similarity, the MoI were associated with a robust repulsion effect wherein CA3/DG representations of the overlapping routes were *less similar* than representations of non-overlapping routes. Importantly, there was no evidence of a corresponding decrease for similarity scores in visual cortical areas at the MoI; instead, similarity scores in visual cortical areas only decreased *after* the MoI (**Fig. 3B**). In fact, the decrease of similarity scores in visual cortical areas after the MoI was a mirror-image of CA3/DG, where similarity scores *increased* after the MoI. Thus, these data establish that CA3/DG repulsion occurred (a) precisely when internal beliefs became distinct and (b) before there was any hint of differentiation within visual cortical areas.

Our second approach involved *manipulating* participants’ internal beliefs while controlling for visual input. Specifically, during fMRI scanning, each trial was preceded by a probabilistic cue that was intended to influence participants’ internal beliefs about the route’s destination. This allowed us to compare fMRI pattern similarity for overlapping routes when cues correctly indicated that the routes were headed to distinct destinations (valid-valid pairs of trials) versus when the cues incorrectly indicated that the routes were headed to the same destination (valid-invalid pairs of trials). Critically, in both cases, we were comparing pairs of route images that were initially identical (same segment) but terminated at distinct destinations. This manipulation revealed that, during the early portion of the route, CA3/DG repulsion depended on the *belief* that routes were headed to distinct destinations (**Fig. X**). In fact, so long as the cues were distinct, repulsion effects were evident in CA3/DG before there were *any differences* in the overlapping route images (i.e., during the same segment; **Fig. X**). The fact that repulsion effects preceded visual differences in the overlapping route images provides even stronger evidence linking repulsion to internal beliefs, as opposed to CA3/DG inheriting or amplifying subtle differences in visual cortical areas.

In addition to establishing that repulsion depends on distinct internal beliefs, our findings strongly reinforce the idea that repulsion also depends on the similarity of visual input into the hippocampus. For example, CA3/DG repulsion selectively occurred early in the trial, when routes were most similar and when similarity scores in visual cortex were highest (**Fig. X**). Even within the similar segment alone, similarity scores in visual cortical areas markedly decreased from the early-similar to late-similar segments while CA3/DG similarity scores markedly increased. In fact, CA3/DG ‘flipped’ from a negative similarity score in the early-similar segment (repulsion) to a positive similarity score in the late-similarity segment. Our interpretation of these data is that the hippocampus plays a critical role in differentiating events when sensory areas fail to do so, but once sensory representations are sufficiently distinct, the hippocampus is no longer needed. This ‘tradeoff’ between the hippocampus and visual cortical areas can also be conceptualized as a shift from internal to external representations. From this perspective, because sensory evidence is ambiguous early in the trial, there is greater reliance on (competing) internal representations supported by the hippocampus (namely, predictions about where the route will go)^31^. Later in the trial, sensory evidence becomes less ambiguous and predictions are less important, thereby shifting processing toward external representations encoded by visual cortex. In fact, an interesting possibility is that repulsion of internal representations within the hippocampus early in the trial directly facilitates external visual attention later in the trial^8^.

While the phenomenon of repulsion is not accounted for or predicted by many of the leading theories of memory and hippocampal function^1,3,4,15^, the *non-monotonic plasticity hypothesis* has specifically been developed to account for repulsion-like effects in behavior and the brain^13,32^. According to this theory, repulsion reflects long-term plasticity that specifically occurs when activation of a target memory spreads to similar, competing memories. Central to this theory is the idea that when a competing memory is moderately activated, it is subject to synaptic weakening that reduces its connections to the target^16,33,34^. In the extreme, selective pruning of target-competitor connections can result in target-competitor similarity that is *lower* than similarity between unrelated events^16,32^— i.e., a repulsion effect. These changes are thought to be adaptive in that they reduce the potential for memory interference in a very targeted manner^16,35–37^. Our findings strongly align with core predictions from this theory. That said, one of the most striking aspects of the current findings is that repulsion effects were highly dynamic and transient within a trial. This raises the question of whether the effects we observed necessarily required, or are best explained by, long-term plasticity.

An alternative possibility is that repulsion reflects dynamic lateral inhibition, without the need for long-term plasticity. From this perspective, activation of neurons associated with a target memory immediately suppresses neurons associated with the competitor. However, this account would require that similar events activate cells that feed into common inhibitory interneurons. In this case, stronger activation of a target would directly inhibit the competitor via the shared inhibitory interneurons. One of the appeals of a lateral inhibition account is that it can explain why the repulsion effects tended to ‘peak’ at particular points in time. Namely, we observed a peak in the repulsion effect immediately after the cue and then again at the moment of insight (**Fig. X**). In other words, whenever the target activation was (putatively) boosted, this may have increased lateral inhibition of the competitor. That said, while lateral inhibition is a core feature of winner-take-all models that explain competitive dynamics in the hippocampus^38,39^, these models have not explicitly referenced or attempted to explain the phenomenon of repulsion.

The fact that we observed repulsion effects in CA3/DG, but not CA1, is consistent with evidence from human^9,11,14,15,19,40^ and rodent studies^1–3,41^ which have consistently found that the CA3-dentate gyrus circuit is involved in disambiguating overlapping events. However, an interesting question is whether CA3 and dentate gyrus differentially contribute to the observed repulsion effects? Although our scanning protocol did not allow us to confidently separate CA3 from dentate gyrus, evidence from rodent studies suggests potential distinctions between these subregions^42,43^. Whereas dentate gyrus is thought to perform relatively automatic pattern separation owing to sparse coding^44^ and strong lateral inhibition^38,39^, CA3 exhibits attractor dynamics that allow for flexible changes between pattern completion and pattern separation processes^3,43,45^. CA3 representations are also thought to be less strictly dictated by sensory input. For example, CA3 supports recall of past experience^46^, anticipation of future experience^47^, and the interpretation of ambiguous sensory input^48^. Thus, our findings in CA3/DG align with many of the functions that have been individually ascribed to these subregions. With more targeted fMRI protocols, it may be possible to tease apart and functionally dissociate the contributions of these subregions to repulsion effects in humans^13,18^.

Taken together, our findings demonstrate that representations of overlapping spatial routes in human CA3 / dentate gyrus exhibit dramatic repulsion effects when visual input is extremely similar (or even identical), but internal beliefs are distinct. By linking repulsion effects to internal beliefs, we provide insight into *when* and *why* hippocampal repulsion occurs. These findings have implications for theories of hippocampal function and are broadly relevant to the fields of human episodic memory and rodent spatial navigation.

## METHODS

### Participants

A target sample size of 40 participants was determined in advance. To reach this sample size, forty-eight participants (27 female; mean age = 20.40 years, range = 18–32) were enrolled following procedures approved by the University of Oregon Institutional Review Board. Written informed consent was collected for each participant prior to the experiment. All participants were right-handed, native-English speakers with normal or corrected-to-normal vision, with no self-reported psychiatric or neurological disease. Three participants were excluded because they ended the experiment early (n = 2) or exited the scanner in the middle of the experiment (n = 1). Five participants were excluded because they failed to reach a pre-determined behavioral threshold (see ‘*Post-test 1,’* below, for details). All participants received monetary compensation for participating.

### Stimuli

The stimuli consisted of eight routes, each corresponding to a stream of 100 images depicting a ‘walk’ through the University of Oregon campus. Images were screenshots taken from videos at a constant time interval. The videos were recorded from an egocentric perspective while a researcher walked along the route. All routes started at the same 4-way intersection on campus and ended at eight distinct destinations which were named after a visual object at the destination (e.g., ‘pole’ or ‘window’). The 8 routes comprised 4 pairs of overlapping routes (**Fig. 1A**). Each pair of overlapping routes left the starting intersection at a different cardinal direction (north, south, east, west). For each overlapping route pair, the first 25 images of the route were identical to each other. Thus, during this initial segment (‘same segment’), it was impossible to distinguish overlapping routes from each other. The next 50 images were extremely similar, but not identical, across the overlapping routes (‘similar’ segment). Specifically, the images were taken from videos that travelled along the same path, but were recorded separately. Thus, images might differ in terms of people, bicycles, shadows, etc. For the last 25 images (‘different segment’), the overlapping routes diverged (each route turned in an opposite direction) before reaching their respective destinations (**Fig. 1A-B**). Each participant studied 2 pairs of routes (i.e., 4 routes). The assignment of routes was counterbalanced across participants such that each subject either studied the north/south routes or the east/west routes.

### Experimental procedure

#### Overview

The first part of the experiment took place in a testing room outside of the MRI scanner. In the testing room, informed consent was obtained and participants were given instructions for the full experiment. Then, participants completed a Learning phase (studying the routes) before entering the MRI scanner. Inside the scanner, participants first completed 2 rounds of a Practice phase (not scanned), which combined additional study and a Pre-Test. Then, participants completed 10 rounds of a Cue phase (scanned), which was the main experimental task. After exiting the scanner, participants completed two Post-Tests that assessed memory for the routes. The experiment was implemented in PsychoPy2021.2.3 and lasted ∼3 hours, with ∼2 hours inside the scanner.

#### Study Phase

During the Study phase, participants viewed each route 4 times in random order. On each trial, the 100 images for a given route were shown in succession. Each image appeared for 240 ms and was immediately followed by the next image. After all images from the route were shown (24 seconds), the destination for the route appeared (2000 ms). The destination was followed by a white fixation cross (3000 ms) and then the next trial started. No behavioral responses were required during this phase. Participants were instructed to pay careful attention to each route so that they would late be able to predict each route’s destination.

#### Pre-test Phase

During each round of the Pre-test phase (2 rounds, total), participants again viewed each route 4 times in random order. The timing of the image presentation in the Pre-test phase—and in all subsequent phases—was identical to the Study phase (i.e., 240 ms per image). For 3 of the 4 presentations, the trial was identical to the Study phase. However, the remaining presentation functioned as a test trial. On these test trials—which were unpredictable from the participants’ perspective—the route paused once per segment (three pauses total per trial). At each pause, participants were shown all 4 possible destinations (distributed in a single row). The destination was only represented by a text label (e.g., ‘pole’). Participants had a maximum of 4000 ms to select the correct destination for the current route by pressing one of four keys on a button box held in their right hand. If they answered within the allotted time, the words ‘sure’ and ‘unsure’ then appeared on the screen and participants had another 3000 ms (maximum) to respond using the button box to indicate their level of confidence. If participants did not respond within the allotted time for the initial set of four destinations, then the confidence decision was omitted. After the third pause/test, the trial continued to the destination. Pauses were restricted such that they only occurred at picture indices 10-25 for the same segment, 26-75 for the similar segment, and 76-90 for the different segment. Within these ranges, the actual pause was randomly determined on each trial. During the Pre-test phase (and the subsequent fMRI phase), stimuli were presented on a gray background, projected from the back of the scanner. Lights were turned off in the scanner room to ensure better contrast for the display.

#### fMRI Phase

In each round of the fMRI phase (10 rounds, total), participants viewed each route 4 times. Importantly, each trial was preceded by a text cue (1 s) indicating the likely destination (e.g., ‘pole’). Cues were either *valid* (indicating the correct destination) or *invalid* (in which case the cue indicated the overlapping route’s destination). Within each round of the fMRI phase, each route was preceded by a valid cue three times and by an invalid cue once. Thus, cues were 75% valid. However, one of the three valid trials served as a catch trial. Catch trials were identical in procedure and timing to the test trials in the Pre-test phase with the exceptions that catch trials (1) always paused (and only paused once) during the similar segment, (2) ended after the confidence rating (or after the destination selection timed out), (3) were preceded by a cue (which, again, was always valid), and (4) had slightly shorter response windows than Pre-Test trials (maximum of 3000 ms to select the destination and, if applicable, a maximum of 2000 ms for the confidence rating). Non-catch trials were always followed by a white fixation cross for 3000 ms before the start of the next trial. Catch trials were followed by a white fixation cross for 3000 ms + any time that was ‘unused’ for the destination selection and confidence ratings. In other words, the duration of the destination selection + confidence rating (if applicable) + fixation cross always summed to 8000 ms.

For catch trials, the pauses occurred randomly between picture indices 25 and 75, but with the constraint that the four catch trials within a given round were randomly divided into two yoked pairs such that the picture indices within each pair summed to 100. For example, if one catch trial was randomly determined to pause at picture index 65, then the yoked catch trial would pause at picture index 35. This constraint ensured that each fMRI scan was identical in length while still maintaining unpredictability about when routes would pause. Performance on the catch trials was not of interest given that, by design, the trials were always preceded by valid cues. Catch trials were only intended to promote vigilance and to reinforce the validity of the cues.

#### Post-test 1

The first Post-test provided an explicit measure of participants’ ability to discriminate between the overlapping routes (in the absence of any cues). Each trial was similar to the test trials from the Pre-test phase. Here, however, every trial contained four pauses (destination tests) that only occurred during the Similar Segment. The pauses occurred at picture indices 30, 45, 60, and 75. For participants 1–32, each route was tested twice in a random order; for participants 33-40, each route was tested 4 times (the difference across participants was due to a technical error in the computer script). An additional difference relative to the test trials from the Pre-test phase is that when pauses occurred during Post-test 1, participants saw a single display that only presented two destination options: the correct destination (target) and the destination of the overlapping route (competitor). Specifically, participants were given five response options, arranged in a single row, corresponding to “Definitely [Destination X]”, “Probably [Destination X]”, “Unsure”, “Probably [Destination Y]”, and “Definitely [Destination Y].” Participants used the trackpad on the laptop to select a response. There was no time limit for responses during this phase. After the pause/test at picture index 75, the trial ended. After a white fixation cross (3000 ms), the next trial started. Participants who were below 100% accuracy (regardless of confidence) at picture index 75 were excluded from the experiment (n = 5). The rationale for excluding these participants is that they did not demonstrate an ability to consistently differentiate the overlapping routes by the end of the similar segment (even after extensive training).

#### Post-test 2

The second Post-test allowed participants to freely indicate the specific timepoint at which they were confident of the route’s destination. As in Post-test 1, routes appeared 4 times each in random order (again without cues). Each trial started with the presentation of the first image (picture index 1) followed by subsequent images up until participants made a response (using the keyboard) to indicate that they were ‘90% sure’ of the route’s destination. Upon making a button press, the route image was replaced by the four destination labels. Participants used the keyboard to select the destination for the route. After a response was made, a white fixation cross appeared for 3000 ms and then the next trial began. There was no time limit for responses during this phase.

### MRI acquisition

All images were acquired on a Siemens 3T Prisma MRI system in the Lewis Center for Neuroimaging at the University of Oregon. Functional data were acquired with a T2*-weighted echo-planar imaging sequence with partial brain coverage that prioritized full coverage of the hippocampus and early visual cortex (repetition time = 1000 ms, echo time = 33 ms, flip angle = 55°, 66 slices, 1.7 × 1.7 × 1.7 mm voxels). A total of 10 functional scans were acquired. Each functional scan comprised 458 volumes and included 6 s of lead-in time and 6 s of lead-out time at the beginning and end of each scan, respectively. Anatomical scans included a whole-brain high-resolution T1-weighted magnetization prepared rapid acquisition gradient-echo anatomical volume (1 × 1 × 1 mm voxels) and a high-resolution (coronal direction) T2-weighted scan (0.43 × 0.43 × 1.8 mm voxels) to facilitate segmentation of hippocampal subfields.

### Anatomical data preprocessing

Preprocessing was performed *using fMRIPrep* 21.0.1 (RRID:SCR_016216), which is based on *Nipype* 1.6.1 (RRID:SCR_002502). The T1-weighted (T1w) image was corrected for intensity nonuniformity (INU) with N4BiasFieldCorrection54 (ANTs 2.3.3, RRID: SCR_004757), and used as the T1w reference throughout the workflow. The T1w reference was skull-stripped with the antsBrainExtraction.sh workflow (ANTs) in Nipype, using OASIS30ANTs as the target template. Brain tissue segmentation of cerebrospinal fluid (CSF), white-matter (WM), and gray-matter (GM) was performed on the brain-extracted T1w using FAST (FSL 6.0.5.1:57b01774, RRID:SCR_002823). Brain surfaces were reconstructed using recon-all (FreeSurfer 6.0.1, RRID:SCR_001847). Volume-based spatial normalization to one standard space (MNI152NLin2009cAsym) was performed through nonlinear registration with antsRegistration (ANTs 2.3.3), using brain-extracted versions of both T1w reference and the T1w template. *ICBM 152 Nonlinear Asymmetrical template version 2009c* was selected for spatial normalization (RRID:SCR_008796; TemplateFlow ID: MNI152NLin2009cAsym).

### Functional data preprocessing

For each participant, a reference volume and its skullstripped version were generated for each of the 10 scan runs by aligning and averaging a single-band reference. For each participant, a fieldmap was collected and estimated based on two echo-planar imaging (EPI) references with topup (FSL 6.0.5.1:57b01774). The estimated fieldmap was then aligned with rigid-registration to the target EPI reference run and the field coefficients were mapped on to the reference EPI. Each scan run was slice-time corrected to 0.445s (0.5 of slice acquisition range 0s-0.89s) using 3dTshift from AFNI (RRID:SCR_005927). The single-band reference was then co-registered to the T1w reference using bbregister (FreeSurfer). Several potentially confounding variables were computed, including: framewise displacement (FD), DVARS, and three region-wise global signals. Additionally, a set of physiological regressors were extracted to allow for component-based noise correction (CompCor). Principal components were estimated after high-pass filtering the preprocessed BOLD time-series (using a discrete cosine filter with 128 s cut-off) for the anatomical CompCor variants (aCompCor). Frames that exceeded a threshold of 0.5 mm FD or 1.5 standardized DVARS were annotated as motion outliers.

The first 6 volumes of each scan run (lead-in time) were discarded. Then, 10 brain masks were generated by fMRIPrep for each of the 10 functional scans. The intersection of all 10 masks was used to perform brain extraction. For each scan run, each voxel was scaled at a mean equal to 100, with an upper bound of 200 and a lower bound of 0. A high-pass filter of 128 seconds was applied to each scan run. Separate categorical regressors were generated to indicate volumes more than 3 standard deviations above or below the global mean or volumes with FD higher than 0.5 mm. To control for nuisance variables, for each scan run a GLM was then applied that included these two categorical regressors along with FD, xyz translation, xyz rotation, aCompCor00-05, and the mean CSF value. Finally, in order to reduce noise in the timeseries data, temporal smoothing was applied within each scan such that, for each voxel, the BOLD response at each TR (volume *n*) was replaced by the average of the response at volumes *n*-1, *n*, and *n*+1.

### Regions of interest

A region of interest (ROI) for early visual cortex (EVC) was created from the probabilistic maps of Visual Topography63^49^ in MNI space with a 0.5 threshold. This ROI was transformed into each participant’s native space using inverse T1w-to-MNI nonlinear transformation. For each participant, an ROI for PPA was created by first using Neurosynth^50^ to perform a meta-analysis with the keyword “place”. Results of the meta-analysis were thresholded by a z-score > 2 using the “associative test” option in Neurosynth. We visually inspected the whole-brain results and manually selected the two largest clusters that were spatially consistent with PPA. One cluster was in the right hemisphere (247 voxels) and one cluster was in the left hemisphere (163 voxels). These clusters were combined into a single PPA mask. This mask was then transformed into each participant’s native space using the inverse T1w-to-MNI transformation. To create hippocampal ROIs, we used the Automatic Segmentation of Hippocampal Subfields (ASHS) 64 toolbox^51^ with the upenn2017 atlas. This generated subfield ROIs in each participant’s hippocampal body, including CA3/DG (which included CA2, CA3, and dentate gyrus) and CA1. The most anterior and posterior slices of the hippocampal body were manually determined for each participant based on the T2-weighted anatomical image. Each participant’s subfield segmentations were also manually inspected to ensure the accuracy of the segmentation protocol. Then, each subfield ROI was transformed from the T2 space into the T1 space using the T2-to-T1w transformation, calculated with FLIRT (fsl) with six degrees of freedom, implemented with Nipype. All ROIs were again visually inspected following the transformation to T1 space to ensure the ROIs were anatomically correct.

### fMRI measures of route similarity

fMRI pattern similarity analyses were used to compute the degree of similarity between overlapping routes relative to non-overlapping routes. To account for hemodynamic response lag, the temporally-smoothed fMRI timeseries for each voxel was first shifted by 6 s, such that the first *timepoint* within each trial was defined as the 6^th^ volume relative to the onset of the route images (each TR = 1 s). Pattern similarity analyses were then performed, separately for each of the 24 timepoints, by computing Pearson correlations between trials from *different scan runs* (correlations were never performed within a scan run). For example, timepoint 1 from trial 1 in scan 1 would be correlated with timepoint 1 from trial 1 in scan 2, timepoint 1 from trial 2 in scan 2, all the way through timepoint 1 from the last trial in scan 10. Catch trials were excluded from pattern similarity analyses. All pattern similarity analyses were performed in participants’ native space and correlation coefficients were Fisher z-transformed before any averaging (hereinafter referred to as pattern similarity).

Of central interest was the similarity between overlapping routes *relative* to the similarity between non-overlapping routes. Specifically, for each participant and each pair of overlapping routes, the mean pattern similarity between non-overlapping routes was subtracted from the mean pattern similarity between the overlapping routes, yielding a *similarity score*. For initial analyses, similarity scores were restricted to trials with valid cues, meaning that pattern similarity was always computed between pairs of trials (whether they were overlapping routes or non-overlapping routes) for which each route was preceded by a valid destination cue (‘valid-valid’ similarity scores).

For subsequent analyses, in order to test for an influence of cue validity, we separately computed ‘valid-invalid’ similarity scores. As with valid-valid similarity scores, the valid-invalid similarity scores were computed by subtracting mean pattern similarity between non-overlapping routes from the mean pattern similarity between overlapping routes. However, valid-invalid similarity scores were based on pattern similarity between pairs of trials (whether they were overlapping routes of non-overlapping routes) for which one route was preceded by a valid cue and the other route was preceded by an invalid cue. It is important to emphasize that, in terms of the actual *route images* that participants saw there was no distinction between valid-valid overlapping routes versus valid-invalid overlapping routes. In other words, whether route 1 was preceded by a valid cue or an invalid cue, and whether route 2 was preceded by a valid cue or an invalid cue had no bearing on the route images participants saw—these overlapping routes always had same, similar and different segments and terminated at distinct destinations. However, in the specific case when one of the cues was valid *and* the other cue was invalid, then participants were led to believe that the two routes were headed to the same destination. For example, if the route 1 cue was valid and the route 2 cue was invalid, then both trials would be preceded by a cue indicating the route 1 destination (because invalid cues always referred to the destination of the overlapping route). Pattern similarity between pairs of invalidly-cued trials (invalid-invalid similarity) was not included in any of the similarity scores reported.

Although similarity scores were always computed separately for each timepoint in a route (24 timepoints per route), we report many of the analyses as a function of route segment (same, early-similar, late-similar, different) instead of route timepoint. In these cases, we simply averaged similarity scores across the timepoints within each segment (e.g., an average of the 6 similarity scores within the same segment).

### Moment of Insight (MoI) analysis

Behavioral responses from Post-test 2 were used to identify the specific point in time at which participants were confident (‘90% sure’) about each route’s destination. We refer to this as the ‘moment of insight’ (MoI). For each participant, each route was tested 4 times in Post-test 2. For each pair of overlapping routes (8 total trials), we pooled all responses and defined the MoI as the range of these response (minimum to maximum, rounded to nearest whole numbers). The rationale for pooling across overlapping routes (as opposed to generating a separate MoI for each route) was that fMRI similarity scores for overlapping routes necessarily reflected the relative similarity or a *pair* of overlapping routes. Thus, it was preferable for the MoI to also be at the level of overlapping route pairs. All timepoints prior to the MoI are referred to as pre-MoI and all timepoints after the MoI are referred to as post-MoI. As with other analyses that were based on route segments (see above), similarity scores were first computed at individual timepoints and then similarity scores were averaged across all timepoints within the pre-MoI, MoI and post-MoI segments.

## Acknowledgements

This work was supported by NIH-NINDS 2R01NS089729 to B.A.K. and by NIH-NINDS F31NS126016 to G.W.

## Author Contributions

S.H., G.W., and B.A.K. designed the experiment. S.H. and G.W recruited participants and collected data. G.W. and B.A.K. analyzed the data and wrote the manuscript.

## Conflicts of Interest

The authors declare no conflicts of interest.

## Notes

### Competing Interest Statement

The authors have declared no competing interest.

