## Supplementary Tables for "Repulsion of CA3 / dentate gyrus representations is driven by distinct internal beliefs in the face of ambiguous sensory input"

**Supplementary Table 1**. Behavioral performance on the Pre-Test. Data represent the mean percentage of total trials associated with each response type ± the standard deviation across participants.

| | **All responses (regardless of confidence)** | | | **High confidence responses only** | | |
|---|---|---|---|---|---|---|
| | Target | Competitor | Non-overlapping | Target | Competitor | Non-overlapping |
| Same segment | 40.00 ± 17.03% | 36.88 ± 15.75% | 5.94 ± 11.67% | 10.12 ± 14.04% | 14.29 ± 12.68% | 2.98 ± 9.61% |
| Similar segment | 73.75 ± 20.57% | 18.44 ± 16.98% | 1.88 ± 4.52% | 58.13 ± 26.49% | 4.38 ± 9.19% | 0.31 ± 1.98% |
| Different segment | 93.13 ± 11.99% | 1.88 ± 4.52% | 3.44 ± 6.93% | 89.38 ± 15.90% | 1.25 ± 3.80% | 1.56 ± 5.05% |

**Supplementary Table 2**. Behavioral performance on Post-test 1. Data represent the mean percentage of total trials associated with each response type ± the standard deviation across participants.

| | **Target** | | | **Competitor** | |
|---|---|---|---|---|---|
| Picture Index | Definitely | Probably | Unsure | Probably | Definitely |
| 30 | 30.16 ± 27.84% | 17.19 ± 18.11% | 42.66 ± 35.69% | 8.44 ± 14.81% | 1.56 ± 4.85% |
| 45 | 82.03 ± 23.05% | 3.59 ± 7.47% | 12.66 ± 20.73% | 1.72 ± 5.48% | 0.00 ± 0.00% |
| 60 | 91.41 ± 15.55% | 5.00 ± 11.25% | 2.97 ± 10.01% | 0.63 ± 2.76% | 0.00 ± 0.00% |
| 75 | 99.38 ± 3.95% | 0.63 ± 3.95% | 0.00 ± 0.00% | 0.00 ± 0.00% | 0.00 ± 0.00% |
